# New insights on the interaction mechanism of rhTNFα with its antagonists Adalimumab and Etanercept

**DOI:** 10.1101/2020.06.21.163824

**Authors:** María Angélica Contreras, Luis Macaya, Pedro Neira, Frank Camacho, Alaín González, Jannel Acosta, Raquel Montesino, Jorge Roberto Toledo, Oliberto Sánchez

## Abstract

TNFα is a pro-inflammatory cytokine that is a therapeutic target for inflammatory autoimmune disorders. Thus, TNFα antagonists are successfully used for the treatment of these disorders. Here, new association patterns of rhTNFα and its antagonists Adalimumab and Etanercept are disclosed. Active rhTNFα was purified by IMAC from the soluble fraction of transformed *E. coli.* Protein detection was assessed by SDS-PAGE and western blot. The K_D_ values for rhTNFα interactions with their antagonists were obtained by non-competitive ELISA and by microscale thermophoresis. Molecular sizes of the complexes were characterized by SEC-HPLC. Surprisingly, both antagonists recognized the monomeric form of rhTNFα under reducing and non-reducing conditions, indicating unexpected bindings of the antagonists to lineal epitopes and to one protomer of rhTNFα. Binding curves of two phases with low and high K_D_ values (<10^−9^ M and >10^−8^ M) were observed during thermophoresis experiments, suggesting the generation of complexes with different stoichiometry, which were confirmed by SEC-HPLC. This pioneer investigation revealed interactions of rhTNFα with Adalimumab and Etanercept never described before, which constitute valuable data for future approaches into the study of their interaction mechanism.

## Introduction

Tumor necrosis factor alpha (TNFα) is a pleiotropic cytokine that promotes inflammation and immune response (1). TNFα is produced by cells of the immune system as a transmembrane protein arranged in homotrimers (2), and a soluble form of TNFα is also generated by proteolytic cleavage (3). Trimeric forms of TNFα interact with the receptors TNF-R1 and TNF-R2, and activate intracellular signaling (4, 5). Different processes involved in inflammation are mediated by TNFα, such as the expression of adhesion molecules in endothelial cells, the recruitment of immune cells to injury sites, the production of other inflammatory cytokines (IL-1, IL-6, IL-17), among others (6).

Although TNFα has an important physiological function in the immune response, it is known that the uncontrolled production or function of this cytokine are related to inflammatory autoimmune diseases, such as rheumatoid arthritis, psoriasis, inflammatory bowel disease, ankylosing spondylitis, and others (7, 8). Thus, TNFα is the main therapeutic target for these diseases, and its inhibition by biopharmaceuticals has been used as a successful treatment against them (6, 7).

Currently, the anti-TNFα biopharmaceuticals in the market are the monoclonal antibodies Infliximab, Adalimumab, and Golimumab. Also, a soluble TNF-R2 receptor named Etanercept, and a pegylated antibody fragment named Certolizumab pegol are available (9). Several researches have determined the interaction of TNFα and its antagonists. The crystalline structures of the complexes TNFα-Infliximab (10), TNFα-Adalimumab (11), TNFα-Golimumab (12) and TNFα-Certolizumab pegol (13) have been resolved. The structure of the complex TNFR2-TNFα was already disclosed (14), which is supposed to be similar to the interaction between Etanercept and TNFα (15, 16). The interaction among TNFα and every antagonist mentioned above are mainly with conformational epitopes. Two protomers of the TNFα homotrimer conform the binding epitopes for Adalimumab and Etanercept, while the epitopes for Infliximab, Golimumab and Certolizumab pegol are composed by residues from one protomer. Moreover, the dissociation constants (K_D_) of these complexes were determined by Surface Plasmon Resonance (SPR) in several reports (17–19), corroborating the high affinity among the TNFα and the biopharmaceuticals.

All TNFα antagonists bind and neutralize the activity of the cytokine (6, 15). However, the biopharmaceuticals have demonstrated distinct effectiveness in the treatment of different inflammatory diseases. These discrepancies have been explained by differences in the binding site and the interaction type among TNFα and its antagonists, and by the effector functions of each molecule (6, 15, 20). The use of other experimental approaches and new techniques could provide novel information about the nature of these bindings. Therefore, the aim of this paper was to study the interactions between TNFα-Adalimumab and TNFα-Etanercept using immunodetection methods, microscale thermophoresis (MST) and size-exclusion chromatography-High performance liquid chromatography (SEC-HPLC). We produced active recombinant human TNFα (rhTNFα) in the *E. coli* strain SHuffle® T7 Express, and the interactions of rhTNFα with Adalimumab and Etanercept were studied. Our main findings were that Adalimumab and Etanercept recognized monomers and lineal epitopes of rhTNFα. Also, the two-phase binding curves observed by microscale thermophoresis and the results of SEC-HPLC confirmed the formation of different molecular weights complexes for Adalimumab-rhTNFα and Etanercept-rhTNFα.

## Results

### Production of active rhTNFα

After expressing the rhTNFα in the *E. coli* strain SHuffle® T7 Express, it was purified by immobilized metal affinity chromatography (IMAC). SDS-PAGE and western blot showed that most of the contaminant proteins were eliminated in the unbounded material and in the wash fractions. Elution showed band of protein around 17-kDa with a 92% purity (Figure 1).

The rhTNFα biological activity was confirmed by an *in vitro* cytotoxicity assay in L929 cells. The cell viability diminished with increasing concentrations of rhTNFα, similar effects were observed for the recombinant human TNFα (Merck, Germany) when it was used as positive control. The values of EC_50_ were 221.8 pg/ml, and 1012 pg/ml for the rhTNFα and for the positive control, respectively (Figure 2A).

To confirm that the reduction of the cell viability caused by the rhTNFα was not due to contaminant proteins, Adalimumab and Etanercept were used to neutralize the cytotoxicity induced by the rhTNFα. The rhTNFα cytotoxicity was totally neutralized with Adalimumab and Etanercept. The Adalimumab EC_50_ was 78.6 ng/ml, and the Etanercept EC_50_ was 71.2 ng/ml (Figure 2B). The cytotoxicity neutralization in L929 cells confirmed that the reduction in the cell viability was only due to the rhTNFα activity.

**Figure 1.**
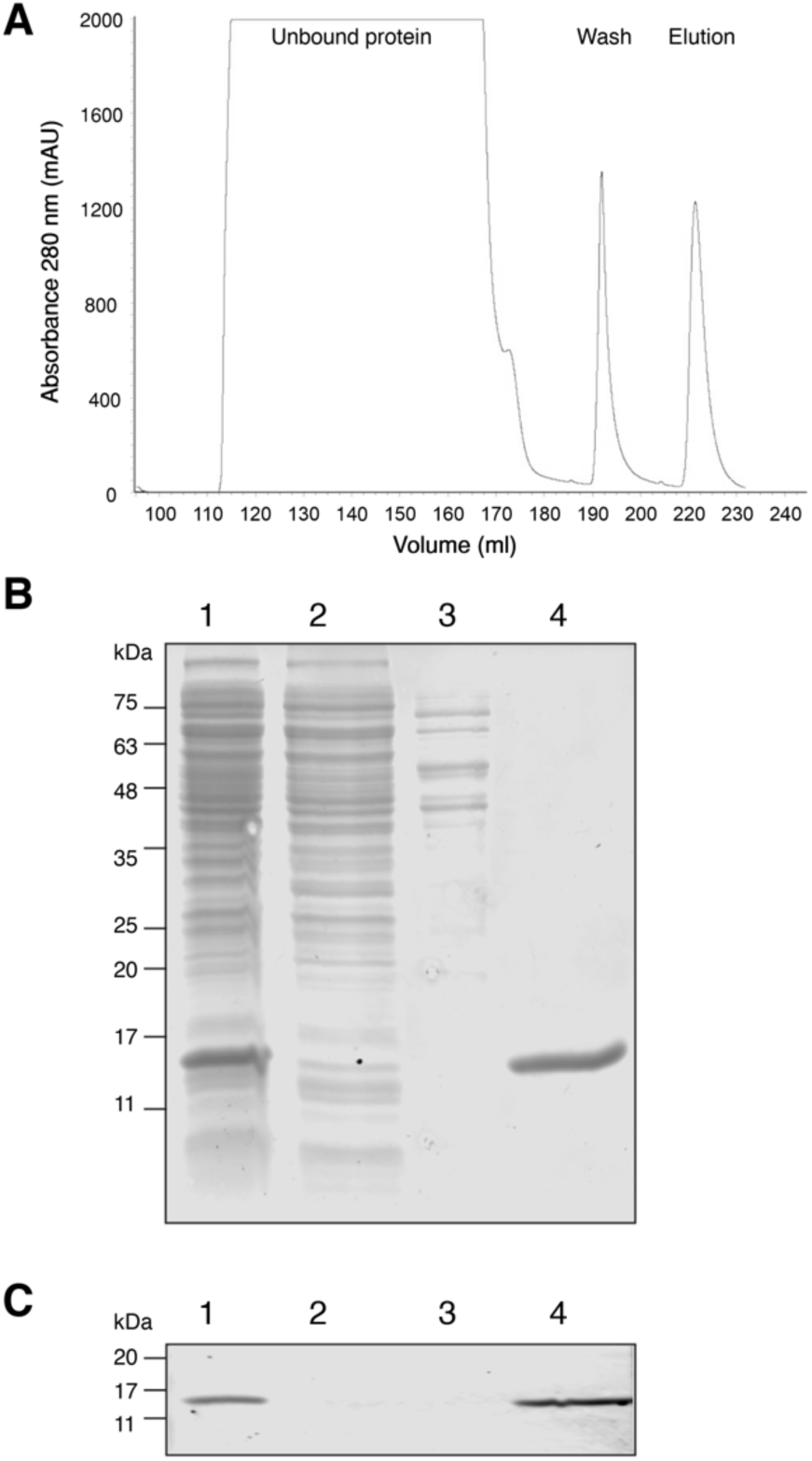
rhTNFα purification from SHuffle® T7 Express *E. coli* by IMAC. (A) Chromatogram of the purification of rhTNFα by Immobilized-metal affinity chromatography (IMAC). (B, C) Analysis of fractions collected in the purification process by 15% SDS-PAGE and Coomassie blue staining (B), and western blot to detect 6xHis tag (C). Lane 1: Input fraction. Lane 2: Flow-through from Ni-NTA affinity column. Lane 3: Wash step. Lane 4: Elution.

**Figure 2.**
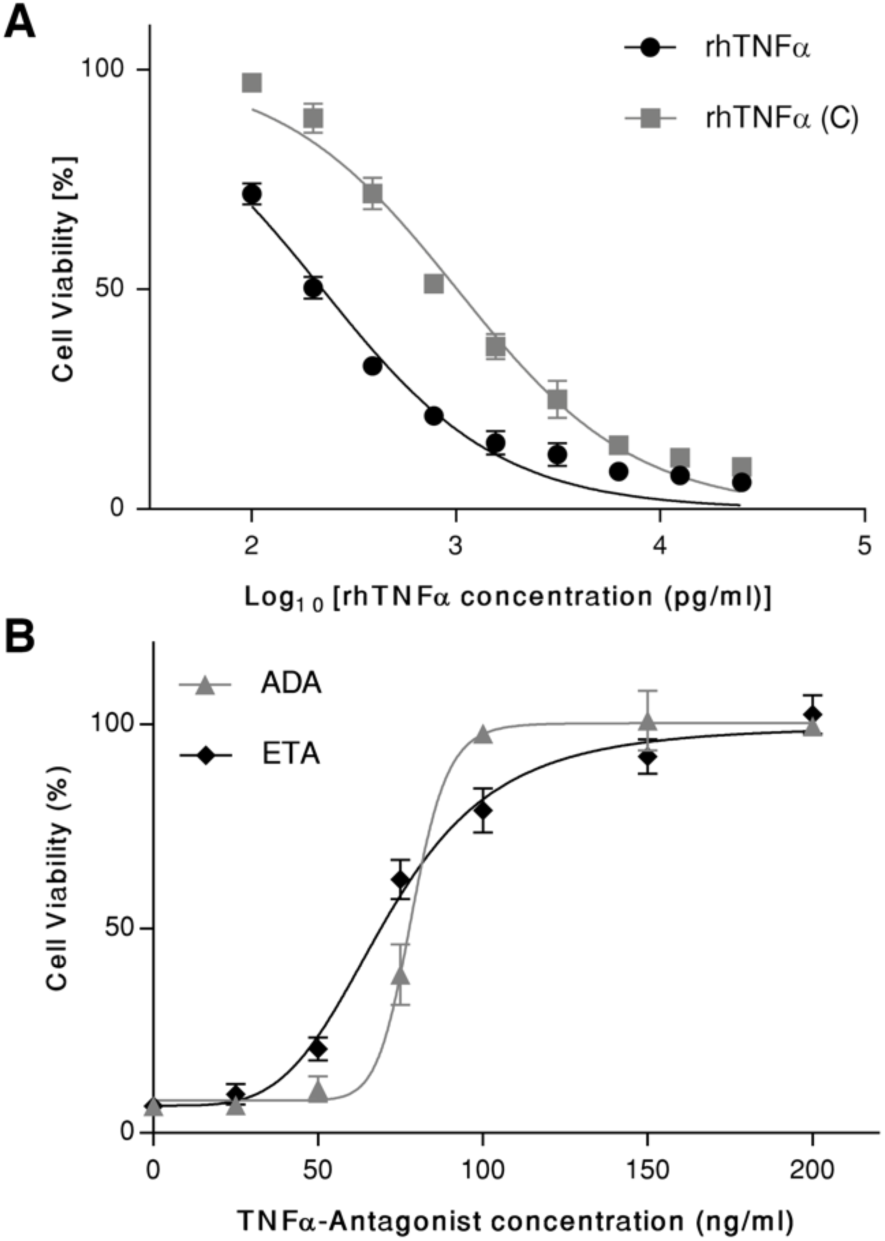
Cytotoxic activity of rhTNFα in L929 cells assay. (A) L929 cells were treated with RPMI-10% FBS medium containing variable concentrations of rhTNFα (0-25 ng/ml). After 20, cells viability was determined by MTT assay. rhTNFα (C) (Merck, Germany) was used as a positive control of activity. The EC_50_ values for rhTNFα (C) and rhTNFα were 1012 pg/ml and 221 pg/ml respectively. (B) L929 cells were treated with RPMI-10% FBS medium containing 25 ng/ml rhTNFα and different concentrations of Etanercept (ETA) or Adalimumab (ADA). After 20 h, cells viability was determined by MTT assay. The EC_50_ values for Adalimumab and Etanercept were 78.6 ng/ml and 71.2 ng/ml respectively. All data were presented as mean ± SD of four replicate wells. Mathematical analysis was performed using Graph Pad Prism 7 software.

### Binding characterization of rhTNFα and its antagonists by immunodetection techniques

The binding among the rhTNFα and its antagonists was detected by native PAGE, SDS-PAGE and western blot in different conditions. The Native PAGE of rhTNFα showed two bands of proteins (Figure 3A), which were recognized by the anti-6xHis Tag antibody in the western blot assay. These bands and an additional one were immuno-identified by Adalimumab and Etanercept (Figure 3B).

**Figure 3.**
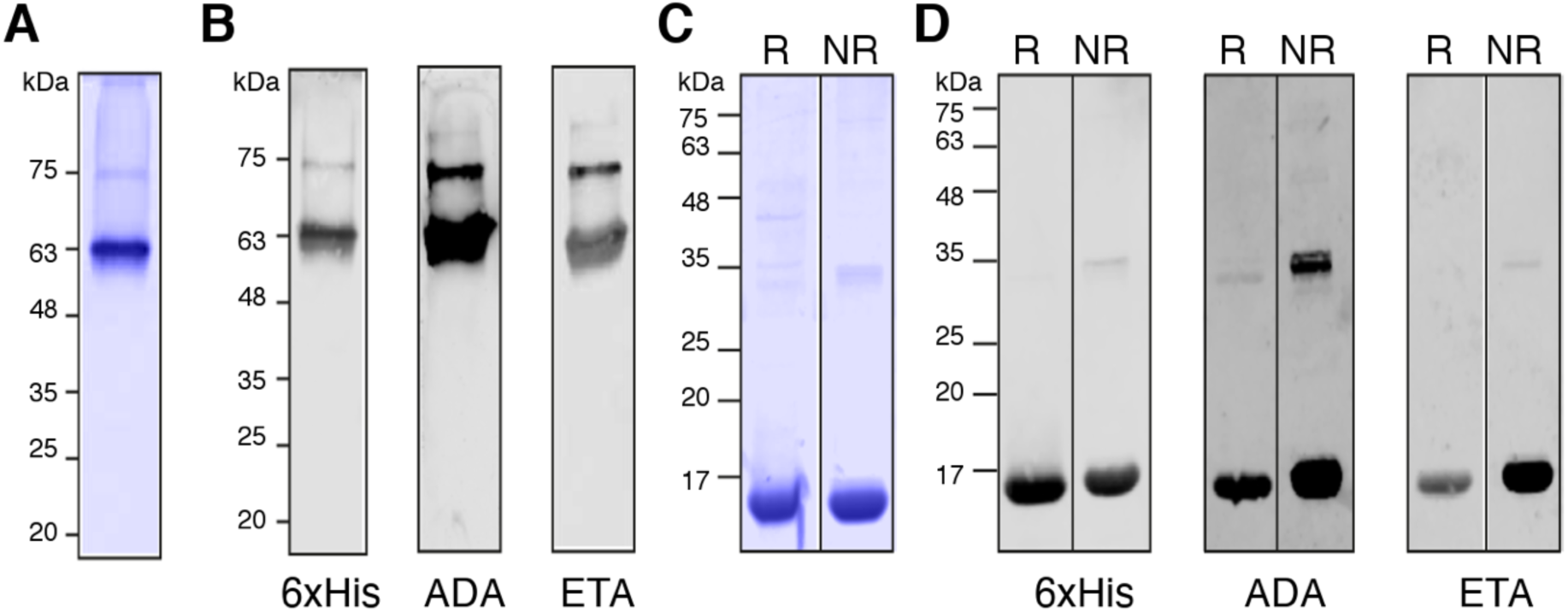
Native and denaturant electrophoresis of rhTNFα and western blot with Adalimumab and Etanercept. (A) Native PAGE and Coomassie blue staining of rhTNFα. (B) Western blot after native PAGE of rhTNFα to detect the protein using anti-6xHis tag antibody (6xHis), Adalimumab (ADA) and Etanercept (ETA). (C) SDS-PAGE and Coomassie blue staining of rhTNFα. (D) Western blot after SDS-PAGE of rhTNFα using anti-6xHis Tag antibody (6xHis), Adalimumab (ADA) and Etanercept (ETA). R: Reducing condition NR: Non-reducing condition.

Also, the rhTNFα was analyzed by SDS-PAGE and western blot under reducing and non-reducing conditions. The SDS-PAGE and the Coomassie blue stain showed different bands and the reinforced band of around 17 kDa corresponded to rhTNFα monomers (Figure 3C). The western blot assays showed that the anti-6xHis antibody, Adalimumab and Etanercept recognized rhTNFα monomers in reducing and non-reducing conditions. Additionally, a band of around 35 kDa was also detected by Adalimumab and Etanercept. It could correspond to dimers of the rhTNFα in non-reducing conditions (Figure 3D).

The Beatty and Beatty approach (21) was used to determine the K_D_ of Adalimumab-rhTNFα and Etanercept-rhTNFα interactions in a non-competitive enzymatic immunoassay. The K_D_ determination of this method require the calculations of the OD_50_ at two concentrations of the rhTNFα. The OD_50_ for Adalimumab at 1,25 and 2,5 µg/ml of rhTNFα were 0.641 nM and 0.622 nM, respectively. The K_D_ for the interaction of TNFα-Adalimumab was 1.32 nM (Figure 4A). For Etanercept-rhTNFα, the OD_50_ using the above concentrations of the rhTNFα were 1.035 nM and 1.072 nM, and the K_D_ for the interaction was 1.99 nM (Figure 4B).

**Figure 4.**
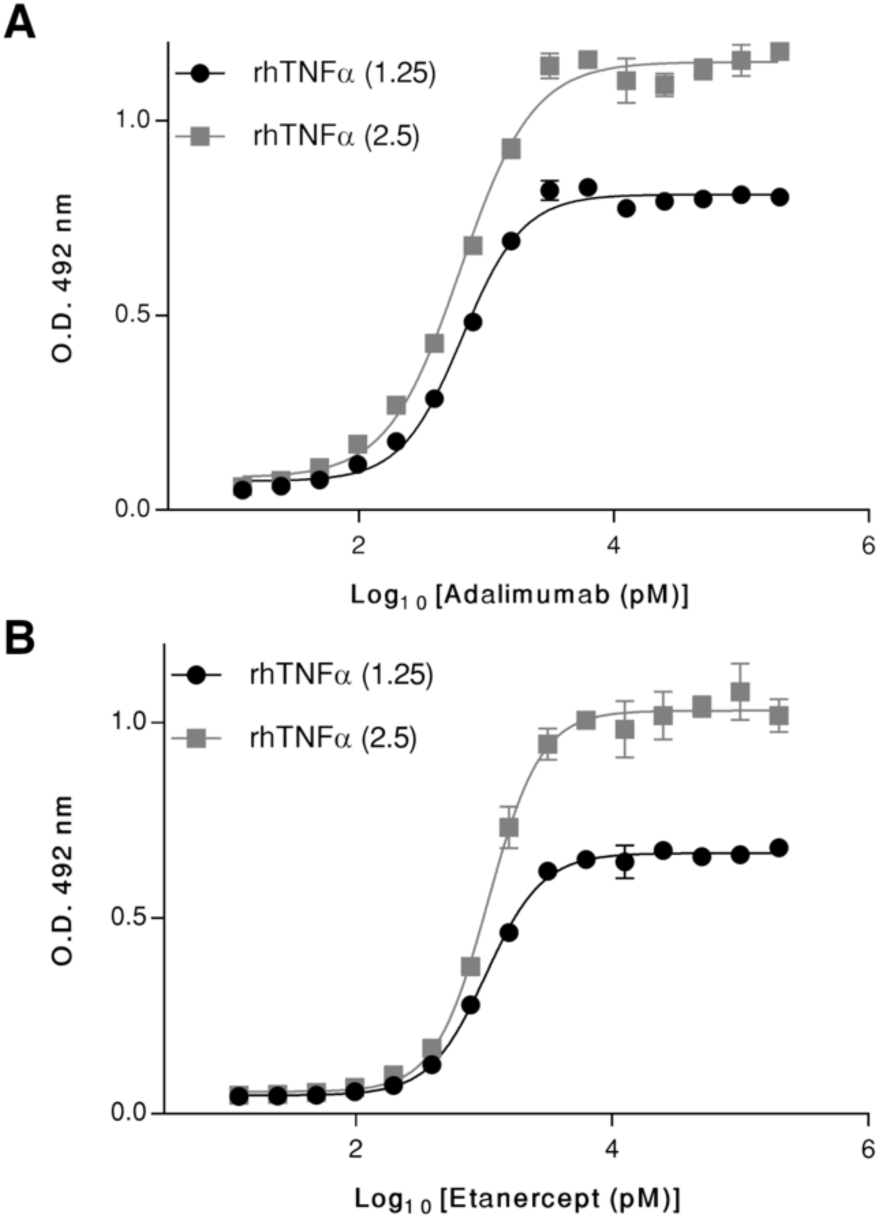
rhTNFα-TNFα antagonists K_D_ determination by a non-competitive enzyme-immunoassay. (A) Experimental binding curve for Adalimumab at two rhTNFα-coating concentrations (1,25 and 2,5 µg/ml), the calculated antibody concentrations at OD_50_ were 641 pM and 622 pM. The K_D_ for the interaction of rhTNFα-Adalimumab was 1.32 nM. (B) Experimental binding curve for Etanercept at two rhTNFα-coating concentrations (1,25 and 2,5 µg/ml), the calculated antibody concentrations at OD_50_ were 1035 pM and 1072 pM. The K_D_ for the interaction of rhTNFα-Etanercept was 1.99 nM. All data were presented as mean ± SD of three replicate wells. Mathematical analysis was performed using Graph Pad Prism 7 software.

### Microscale thermophoresis and SEC-HPLC of rhTNFα and its antagonists

For studying the interaction of rhTNFα-Adalimumab and rhTNFα-Etanercept by MST, it is required the labeling of one molecule in each pair of interaction. As the labeling could alter the ligand binding, two MST experiments was performed for each interaction partners. One by labeling rhTNFα and the other by labeling the antagonist. The rhTNFα and Etanercept were labeled in the amine groups (NHS-coupling). However, Adalimumab was labeled in the thiol groups (Cys-coupling) because any interaction was detected when it was labeled in the amine groups (not shown).

Two MST experiments were performed to study the interaction rhTNFα-Adalimumab. First, the MST was performed using a fluorescent-labeled Adalimumab (Cys-coupling) and rhTNFα without labeling. A binding curve was obtained with a K_D_ value of 29.6 ± 7.4 nM (Figure 5A). A second experiment was performed using the rhTNFα labeled in the amine groups and Adalimumab without labeling. A two-phases equilibrium was observed. At concentrations of Adalimumab from 30.5 pM to 7.8 nM, a K_D_ value of 100.2 ± 187.5 pM was determined, and for Adalimumab concentrations from 1.95 to 1 µM, we obtained a K_D_ value of 21.00 ± 9.43 nM (Figure 5B).

**Figure 5.**
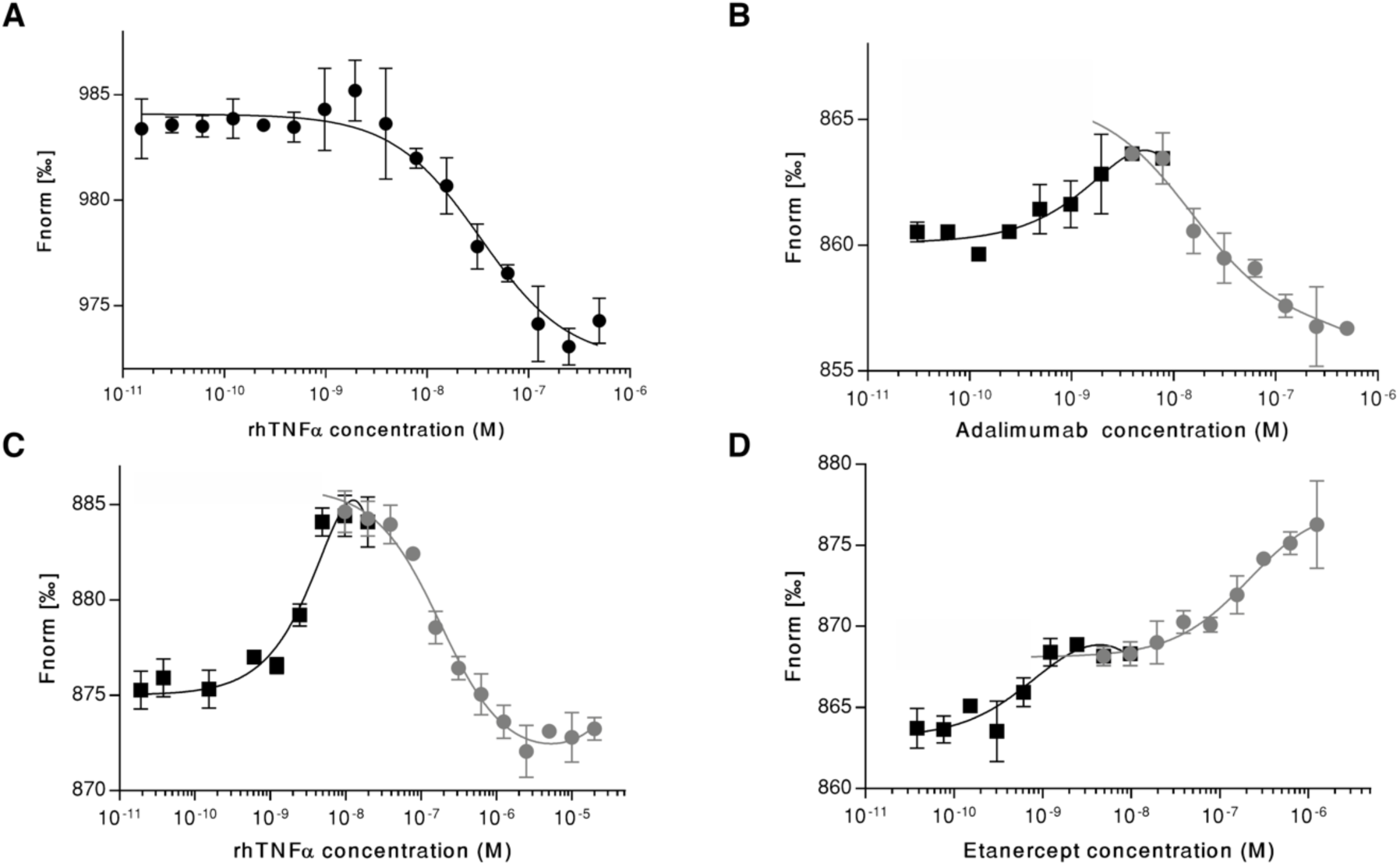
rhTNFα-TNFα antagonists binding studies by microscale thermophoresis. (A) Binding curve of labeled Adalimumab with rhTNFα. Adalimumab (NT-647) was constant 5nM while the concentration of rhTNFα was varied between 500 nM–30.5 pM. The MST traces were analyzed and the K_D_ of the interaction was 29.6 ± 7.4 nM. N=3. (B) Binding curve of labeled rhTNFα with Adalimumab. The MST experiment was performed at constant concentration of labeled rhTNFα (2 nM) and variable concentration of Adalimumab between 30.5 pM and 1 µM. The MST traces were analyzed and two binding curves were detected. The first curve had a K_D_ value of 100.2 ± 187.5 pM, and the second curve had a K_D_ value of 21.00 ± 9.43 nM. N=2. (C) Binding curve of labeled Etanercept with rhTNFα. In the MST experiment, the concentration of labeled Etanercept was constant at 2 nM, while the concentration of the non-labeled rhTNFα was varied between 19.1 pM and 20 µM. The MST traces were analyzed and two binding curves were detected. The first curve had a K_D_ value of 1.25 ± 0.78 nM, and the second curve had a K_D_ value of 175.5 ± 38.2 nM. N=3. (D) Binding curve of labeled rhTNFα with Etanercept. The experiment was performed using labeled TNFα at 2 nM, and the Etanercept concentrations were variable between 38.1 pM and 1.25 µM. The MST traces were analyzed and two binding curves were detected. The first curve had a K_D_ value of 317.8 ± 325.3 pM, and the second curve had a K_D_ value of 183.5 ± 23.4 nM. N=3. All MST experiments were performed using the Monolith NT.115 Pico (NanoTemper Technologies) at 25°C and the results were analyzed in the software M.O. Affinity Analysis v.2.3 (NanoTemper Technologies). The charts were generated in Graph Pad Prism 7 software. All data were presented as mean ± SD of replicates.

The evaluation of the interaction rhTNFα-Etanercept with a similar approach was performed. A two-phase pattern was observed when Etanercept labeled in the amino groups and non-labeled rhTNFα were used. A K_D_ value of 1.25 ± 0.78 nM was obtained at concentrations of the rhTNFα from 19.1 pM to 39.1 nM. At rhTNFα concentrations from 4.88 nM to 20 µM, a K_D_ value of 175.5 ± 38.2 nM was detected (Figure 5C). Finally, the interaction rhTNFα-Etanercept was analyzed using labeled rhTNFα and unlabeled Etanercept. Also, a two-phases equilibrium was detected. The first curve had a K_D_ value of 317.8 ± 325.3 pM, and it involved Etanercept concentrations from 38.1 pM to 39.1 nM. The second curve had a K_D_ value of 183.5 ± 23.4 nM when concentrations of Etanercept from 1.22 nM to 1.25 µM were evaluated (Figure 5D). To corroborate that multiple equilibria detected by MST were due to the formation of complexes with different compositions, samples containing distinct molar ratios of rhTNFα and its antagonists were analyzed by SEC-HPLC. When Adalimumab and rhTNFα were mixed at equimolar concentration, complexes with retention times (RT) between 11 min and 18 min were detected with molecular weight up to 600 kDa, approximately (Figure 6B). When an excess of Adalimumab or rhTNFα were analyzed, we detected the molecule excess and species with variable weights. The Adalimumab excess showed a signal at 17.5 min, which corresponded to complexes of around 650 kDa (Figure 6C). The rhTNFα excess displayed peaks at 17.8 and 19.5 min, which agreed with complexes of molecular weights around 625 and 403 kDa, respectively (Figure 6D).

**Figure 6.**
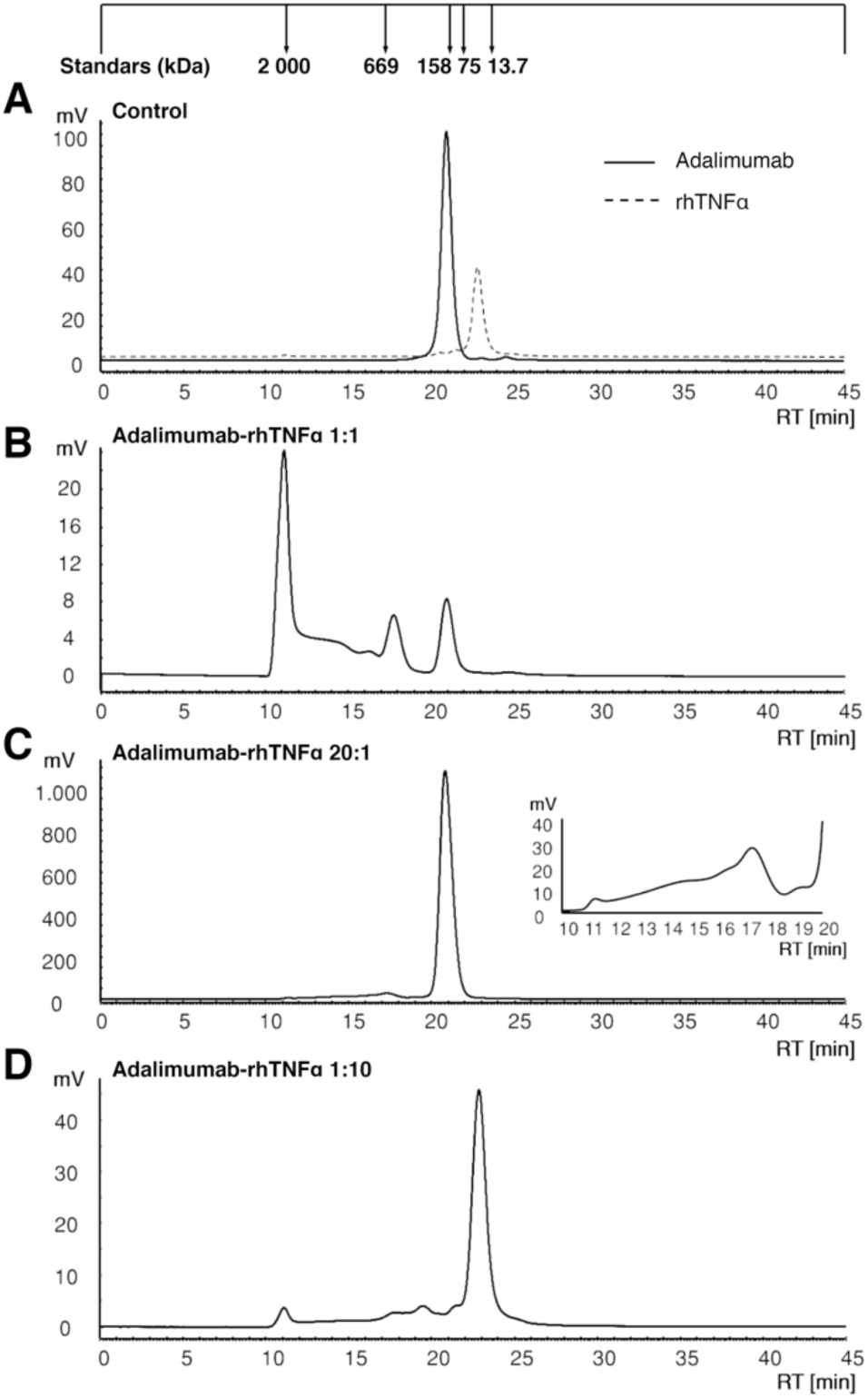
rhTNFα-Adalimumab complexes analysis by SEC-HPLC. (A) Controls. Samples of Adalimumab and rhTNFα at 6.7 µM were analyzed independently. (B) Adalimumab-rhTNFα 1:1. Analysis of a solution containing Adalimumab (6.7 µM) and rhTNFα (6.7 µM). (C) Adalimumab-rhTNFα 20:1. Analysis of a solution containing Adalimumab (670 µM) and rhTNFα (6.7 µM). (D) Adalimumab-rhTNFα 1:10. Analysis of a solution containing Adalimumab (3.35 µM) and rhTNFα (33.5 µM). All separation experiments were performed on a Jasco HPLC system with a TSK Gel G4000SW_XL_ column. A flow rate of 0.5 ml/min and a 50 mM phosphate-300 mM NaCl buffer (pH 7.5) were used for the analysis. The system was calibrated using Blue Dextran 2000 (2.000 kDa), Thyroglobulin (669 kDa), Aldolase (158 kDa), Conalbumin (75 kDa) and Ribonuclease A (13.7 kDa).

The same methodology was used to analyze the interaction between Etanercept and rhTNFα. The Etanercept-rhTNFα interaction at the same concentration showed a wider and asymmetric peak compared to the control, and the signal of rhTNFα was not observed (Figure 7B), meaning that all rhTNFα was bound to Etanercept. When an excess of Etanercept was evaluated, only its signal was detected (Figure 7C), and the rhTNFα excess revealed three peaks at 17.2 min (up to 669 kDa), 19.2 min (around 453 kDa) and 23 min (rhTNFα excess) (Figure 7D).

**Figure 7.**
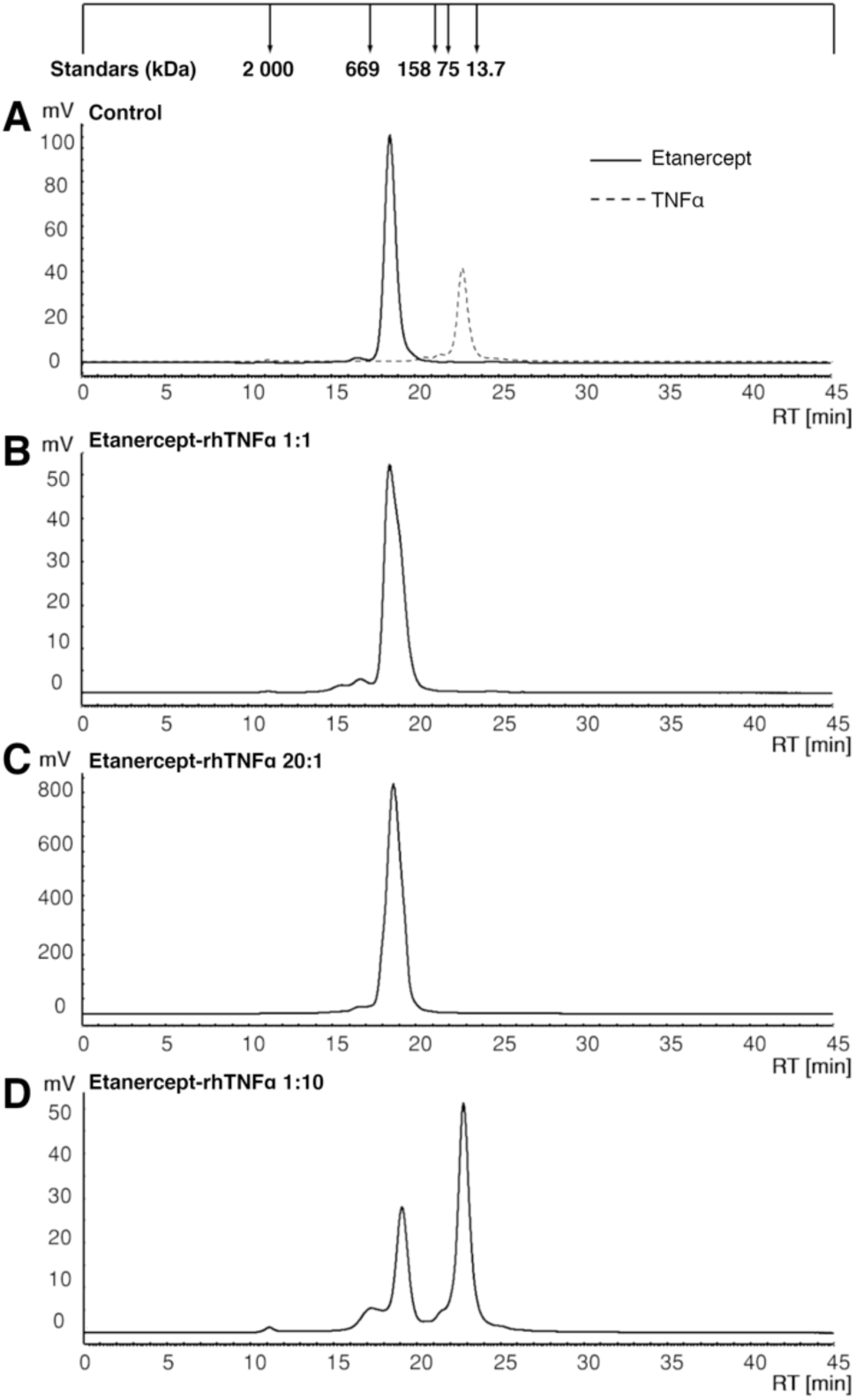
rhTNFα-Etanercept complex analysis by SEC-HPLC. (A) Controls. Samples of Etanercept and rhTNFα at 6.7 µM were analyzed independently. (B) Etanercept-rhTNFα 1:1. Analysis of a solution containing Etanercept (6.7 µM) and rhTNFα (6.7 µM). (C) Etanercept-rhTNFα 20:1. Analysis of a solution containing Etanercept (670 µM) and rhTNFα (6.7 µM). (D) Etanercept-rhTNFα 1:10. Analysis of a solution containing Etanercept (3.35 µM) and rhTNFα (33.5 µM). All separation experiments were performed on a Jasco HPLC system with a TSK Gel G4000SW_XL_ column. A flow rate of 0.5 mL/min and a 50 mM phosphate-300 mM NaCl buffer (pH 7.5) were used for the analysis. The system was calibrated using Blue Dextran 2000 (2.000 kDa), Thyroglobulin (669 kDa), Aldolase (158 kDa), Conalbumin (75 kDa) and Ribonuclease A (13.7 kDa).

## Discussion

TNFα is an important pro-inflammatory cytokine for the immune response. It also has a critical role in inflammatory autoimmune disorders, and the inhibition of TNFα with biopharmaceuticals constitute an effective treatment for them. Adalimumab and Etanercept are the most sold biopharmaceuticals (22), which are two different types of TNFα antagonists. Adalimumab is a human monoclonal antibody (IgG1), and Etanercept is a fusion protein based on the extracellular domain of the human TNF-R2 and the Fc fragment of human IgG1 (6). Here, active rhTNFα was produced in *E. coli*, and its interaction with Adalimumab and Etanercept was studied. The main findings were: 1) Adalimumab and Etanercept recognize rhTNFα monomers, even under denaturant and reducing conditions, suggesting that these molecules could recognize linear epitopes in rhTNFα; 2) At different concentrations of rhTNFα, Adalimumab and Etanercept, complexes with diverse stoichiometry are formed.

Soluble hTNFα is a 51 kDa trimer, which has been described as a necessary condition for its biological activity (5). Here, we produced rhTNFα with similar activity to that reported elsewhere (23, 24). It was used for all experimental procedures.

The interaction between TNFα and biopharmaceuticals has been studied in several reports using different techniques and approaches. The reported crystal structure of the complex TNFα-Adalimumab (PDB code: 3WD5) revealed that the antibody recognized a conformational epitope in TNFα, which is composed by residues from two protomers of the protein (Figure 8A) (11). Considering the structure for the TNFα-TNFR2 complex (PDB code: 3ALQ), it is expected that Etanercept interact with TNFα in the groove formed by two protomers (Figure 8B) (14). However, we observed that Adalimumab and Etanercept recognized rhTNFα monomers in western blot assays after denaturant electrophoresis in reducing and non-reducing conditions. This is the first study that reports the interaction of Adalimumab and Etanercept with TNFα monomers and lineal epitopes.

**Figure 8.**
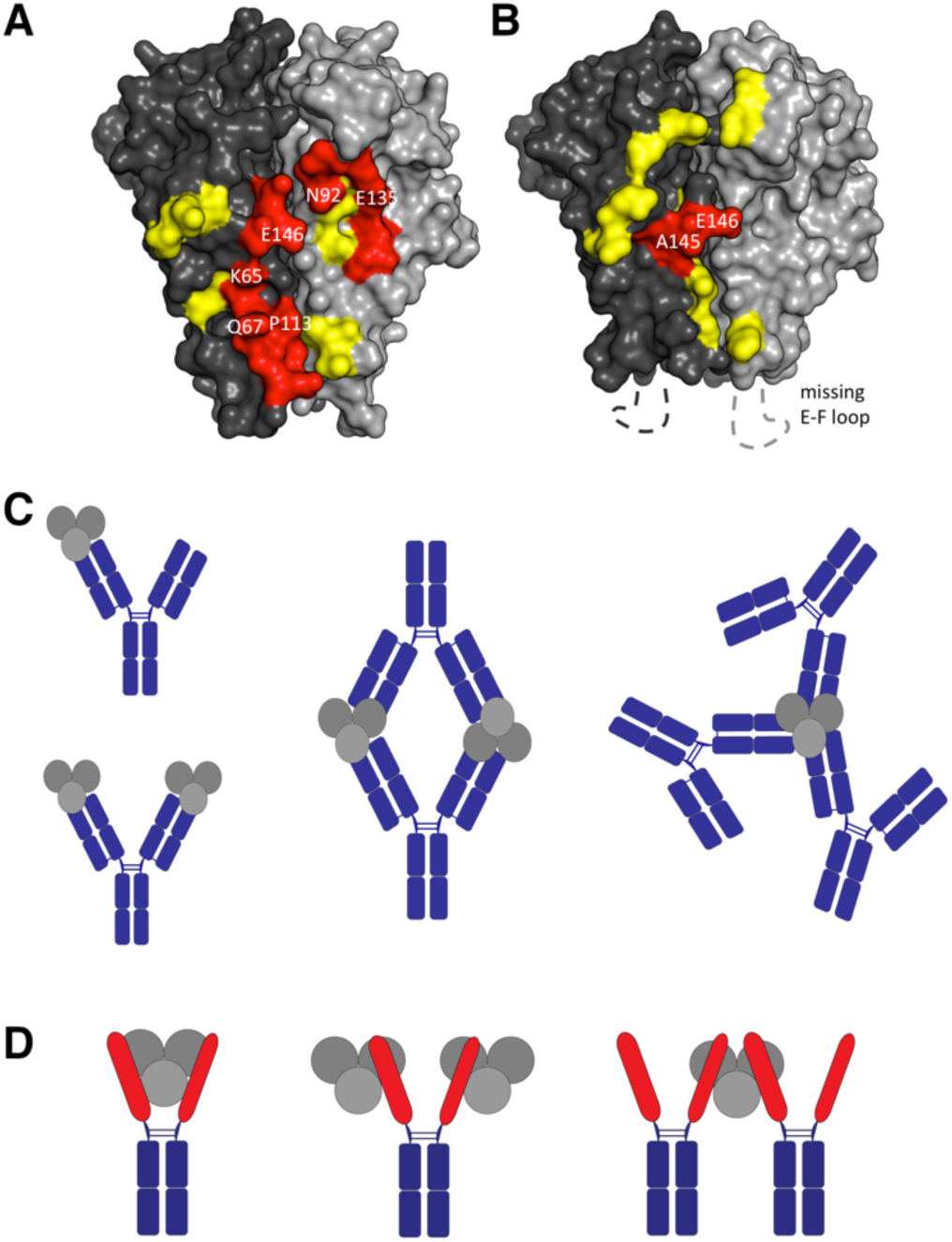
TNFα and TNFα-antagonist interactions. (A, B) Tridimensional representation of TNFα, monomers are presented in different shades of grey and the interfaces of the interaction with Adalimumab (A) and TNFR2 (B) are shown in yellow and red. The red surface are the sequences postulates as lineal epitopes. (C) Illustration of the TNFα-Adalimumab complexes. (D) Illustration of the proposed TNFα-Etanercept complexes.

Analyzing the TNFα-Adalimumab interphase of the crystal structure (11), five linear aminoacidic sequences were detected in TNFα: K_65_-G_66_-Q_67_, K_90_-V_91_-N_92_, E_110_-A_111_-K_112_-P_113_, E_135_-I_136_-N_137_ and A_145_-E_146_-S_147_ (Figure 8A). These sequences could be recognized by Adalimumab in denaturant and reducing conditions. Also, mutations of K_65_, Q_67_, E_135_, and E_146_ by alanine residues generates a 100-fold decrease in the affinity of Adalimumab for its target (11), which confirms the importance of these residues for the TNFα-Adalimumab interaction. On the other hand, the analysis of the TNFα-Etanercept interface showed the sequence D_143_-F_144_-A_145_-E_146_ (Figure 8B), which forms the G-H loop in TNFα and is crucial to the interaction with the Cysteine Rich Domain 2 (CRD2) of TNF-R2 (14). This sequence could be recognized by Etanercept when rhTNFα was run under denaturant and reducing conditions.

As a new approach to study the interactions of rhTNFα with Adalimumab and Etanercept, we used the microscale thermophoresis (MST). This methodology is based on the motion of molecules in microscopic temperature gradients generated in a free fluid. It is an easy and rapid technique, which requires low sample quantities and there are no limitations of molecular weight or size when the interactions are measured. Additionally, the K_D_ of a wide binding variety can be detected in the equilibrium stage by titration experiments (25, 26). Our MST analysis revealed equilibria in two phases. At low molar ratios, we detected low K_D_ values for both complexes (rhTNFα-Adalimumab, rhTNFα-Etanercept) in the first phase of the binding curves, which were consistent with the K_D_ values previously reported (17–19, 27, 28). The second phase of the binding curves obtained at high ligand concentration were unexpected. We postulated that these high K_D_ values corresponded to the formation of complexes with a stoichiometry higher than 1:1 among rhTNFα and its antagonists. Indeed, complexes of high molecular weights were detected by SEC-HPLC. The interaction between rhTNFα and Adalimumab showed complexes higher than 600 kDa. Previous reports have detected the formation of complexes 1:1, 1:2, 2:2, and 3:2 ratios of Adalimumab:TNFα by electron microscopy (29). Moreover, complexes of 4.000 kDa were described by SEC-LS (30). Proposed complexes of Adalimumab-TNFα are presented in the Figure 8C. Additional structures of higher molecular weights could be formed taking into account the multiple sites of interaction available in each molecule, and the flexibility that the hinge region of Adalimumab could provide to the complexes.

Although, the formation of the complex TNFα-Etanercept 1:2 was reported previously when 3-fold Etanercept over TNFα was evaluated by SEC-LS (30), it is generally assumed that one Etanercept bound one TNFα trimer (19, 20). We observed complexes higher than 400 kDa by SEC-HPLC. Unlike the interaction of rhTNFα-Adalimumab, complexes of high molecular weight (sizes up to 2.000 kDa) were not detected for rhTNFα-Etanercept. Considering the results of MST and SEC-HPLC, we hypothesized that complexes TNFα-Etanercept 2:1 might be formed when an excess of TNFα was added, while complexes 1:2 could be formed when there was an excess of Etanercept. Schemes of these complexes are presented in Figure 8D. The rigid structure of Etanercept could prevent the formation of high molecular weight complexes. Our results demonstrated that the binding of rhTNFα with Adalimumab and Etanercept involve the formation of higher structures when an excess of ligand is added. Under these conditions, K_D_ values of rhTNFα-Adalimumab complexes were ten-fold lower than that of rhTNFα-Etanercept complexes. This data could suggest that the assembly of rhTNFα-Adalimumab complexes are favored compared to rhTNFα-Etanercept complexes. It also could be related to the individual structures of Adalimumab and Etanercept.

Under physiological and pathophysiological contexts, TNFα concentrations in serum are around 10 pg/ml (2×10^−12^ M) (31, 32). The Adalimumab and Etanercept concentrations in serum range from 1 to 8 µg/ml (6.7×10^−9^ to 5,3×10^−8^ M) (33–37). The levels of these molecules suggest that the complexes mentioned above could not be formed. However, the accumulation of these molecules in the inflammation sites could favor the formation of high molecular weight complexes. Still, it has not been reported and the relevance of this issue is uncertain.

Overall, novel interaction of rhTNFα with Adalimumab and Etanercept were described. Further studies must be conducted to elucidated if the formation of higher complexes could be relevant for future therapeutic approaches.

## Experimental procedures

### Reagents and antibodies

Chemical compounds were provided by Merck and Sigma-Aldrich. Restriction enzymes were purchased from New England Biolabs. Recombinant human TNFα (GF023) was from Merck. Purified anti-His tag mouse antibody was from BioLegend (USA). Secondary antibodies Alexa Fluor® 680 AffiniPure Donkey anti-mouse IgG and Alexa Fluor® 790 AffiniPure Donkey Anti-human IgG (H+L) were obtained from Jackson ImmunoResearch Inc (USA), and anti-Human IgG (Fc specific) −Peroxidase antibody produced in goat was from Sigma-Aldrich. Two TNFα-antagonists were purchased: Adalimumab (Humira®, Abbie) and Etanercept (Enbrel®, Pfizer).

### Construction of expression vector

The DNA sequence codifying the soluble human TNFα (residues 85-233 from UniProt P01375) was sub-cloned into a pET-22b (+) vector at the XhoI and NdeI sites, generating pET-22b-hTNFα vector. The pET-22b-hTNFα vector permits the TNFα expression under the control of T7 promoter, and it is inducible using isopropyl-β-D-thiogalactopyranoside (IPTG). Furthermore, the gene of TNFα is in frame with a C-terminal His•Tag® to facilitate the detection and purification of recombinant protein.

### Recombinant hTNFα obtention

SHuffle® T7 Express chemically competent *E. coli* (New England Biolabs, England) were transformed with pET-22b-hTNFα vector and selected in LB medium supplemented with 100 µg/ml ampicillin (US Biological, USA). A clone of transformed bacteria was grown in liquid LB-Ampicillin medium at 30°C until O.D. _600 nm_ 0.8; IPTG (US Biological, USA) was added to a final concentration of 0.1 mM. After six hours, bacteria were collected by centrifugation at 6.000 g and resuspended in 50 mM Tris-HCl pH 7.5. Then, cells were disrupted in an Emulsiflex-C5 homogenizer (Avestin, Canada); soluble and insoluble fractions were separated by centrifugation (28960 g, 4°C, 30 min). Immobilized Metal Affinity Chromatography (IMAC) was performed for protein purification using an AKTA Start Chromatography system (GE Healthcare Bio-Science, Sweden). The soluble fraction containing TNFα was diluted (1:1) with equilibrium buffer (50 mM Tris, 5 mM Imidazole, 300 mM NaCl, pH 7.5) and was applied to Ni-charged chelating Sepharose Fast Flow column (GE Healthcare Bio-Science, Sweden). After that, the column was washed with 50 mM Tris-100 mM Imidazole-300 mM NaCl pH 7.5, and the recombinant protein was eluted with 50 mM Tris-300 mM Imidazole-300 mM NaCl pH 7.5. An Amicon Ultra-15 Centrifugal Filter Unit 3kDa (Merck, Germany) was used to concentrate elution fraction containing rhTNFα and exchange the buffer to PBS 1X pH 7.4. The total protein concentration was determined using BCA protein assay kit (Thermo Fisher Scientific, USA). The purity percentage of rhTNFα was calculated by densitometry using freely available ImageJ software (NIH Image, USA).

### Electrophoresis and western blot analysis

For the detection of rhTNFα, 5 µg of protein were loaded on 15% SDS-PAGE, and 8 µg of protein were loaded on 12% native-PAGE. Protein electrophoresis was conducted according to the standard methods (38). The native-PAGE was run using the protocol similar to SDS-PAGE except for the absence of SDS in the loading buffer, the running buffer, and the gel. Gels were stained with Coomassie Blue (Phast Gel ™ Blue R, GE Healthcare, Sweden).

For western blot, proteins were transferred to 0.2 µm nitrocellulose membranes (GE Healthcare Life science, Germany) in a semi-dry transfer system (Bio-Rad, USA). The membranes were blocked with 5% skim milk in TBS for 2 h at room temperature. Primary antibodies (anti-His Tag mouse antibody, Adalimumab and Etanercept) were prepared in TBS containing 2% skim milk, and the membrane was incubated for 2 h with the preparation. Membranes were washed three times and incubated for 1h with secondary antibodies conjugated with a fluorophore (anti-mouse IgG-Alexa Fluor 680 and anti-human IgG-Alexa Fluor 790). The blots were detected and digitalized using the Odyssey infrared imaging system (LI-COR Biosciences, USA).

### L929 cytotoxicity assay

L929 cells (ATCC) were seeded in 96-well plate at a 3×10^4^ cells per well density and were cultured in RPMI-1640 medium (Biological Industries, Israel) containing 10% FBS (Biological Industries, Israel) and 50 µg/ml Neomycin (Sigma-Aldrich, Switzerland) with 5% CO_2_ at 37°C. The cells were incubated for 24 h to allow attachment. Then the culture medium was removed and replaced with RPMI medium containing 1 µg/ml D-Actinomycin (Merck, Germany) and different samples of rhTNFα or rhTNFα incubated with TNFα antagonist. Cells were treated for 20 h, and cell viability was determined by MTT assay (39). To MTT assay, the medium was replaced with 110 µl per well of RPMI containing 0.45 mg/ml MTT (Thermo Fisher Scientific, USA). After 4 h of incubation at 37°C in darkness, supernatants were removed carefully, and 100 µl of isopropanol was added to each well for dissolve formazan crystals. The absorbance was recorded at 570 nm in a Synergy™ HTX Multi-Mode Microplate Reader (BioTek). Results were represented as cells viability percent (%V) calculated as %V= (O.D. test group/O.D. control group) x100. Mathematical analysis was performed using Graph Pad Prism 7 software.

### Enzyme Immunoassay

The wells of a high binding 96-well plate were coated with 100 µl of TNFα at 1.25 and 2.5 µg/ml in coating buffer pH 9.6 at 4°C overnight, and blocked with 200 µl blocking buffer (3% BSA in PBS) at 37°C for 2h. Samples of different concentrations of anti-TNFα molecules Adalimumab and Etanercept (200 nM to 12,2 pM) were applied in wells and incubated at 25°C for 1h. Then, the wells were washed with PBS-0.1% Tween 20 for three times; 100 µl of peroxidase-conjugated anti-human IgG antibody dissolved 1:5.000 in 1% BSA-PBS were added to each well. After 1 h, wells were washed with PBS-0.1% Tween 20, and 100 µl of a solution containing OPD (Santa Cruz Biotechnology Inc.) was added to each well. The reaction was left to develop in the dark and was stopped with 2.5 M H_2_SO_4_. Finally, the absorbance was recorded at 492 nm in a Synergy™ HTX Multi-Mode Microplate Reader (BioTek). Mathematical analysis was performed using Graph Pad Prism 7 software.

### Microscale Thermophoresis

The MST experiments were performed in a Monolith NT.115 Pico (NanoTemper Technologies, Germany) using the Monolith Protein Labeling Kit RED-NHS (Amine Reactive), the Monolith Protein Labeling Kit RED-NHS 2nd Generation (Amine Reactive), the Monolith Protein Labeling Kit RED-Maleimide (Cysteine Reactive), and the Monolith NT.115 Premium Capillaries (NanoTemper Technologies, Germany).

For experiments with labeled rhTNFα (Amine Reactive) and TNFα-antagonists, we have kept the concentration of NT-650 labeled rhTNFα constant (2 nM). In contrast, the concentration of Etanercept was varied between 1.25 µM–38.1 pM and the concentration of Adalimumab was varied between 1 µM–30.5 pM. The assays were performed in PBS containing Tween-20 0.05%, at 40% of excitation power, and 40% of MST power. MST traces were analyzed at 5 seconds. For performing experiments with labeled Adalimumab, a fluorescent label (NT-647) was covalently attached to the protein (Cysteine Reactive). In the experiment, the concentration of labeled Adalimumab was constant (5 nM), while the concentration of rhTNFα was varied between 500 nM–30.5 pM. The assay was performed in the buffer provided by the manufacturer containing Tris-HCl 50 mM pH 7.4, NaCl 150 mM, MgCl_2_ 10 mM, and Tween-20 0.05%. After a short incubation, the samples were loaded into MST NT.115 hydrophilic glass capillaries. The experiment was performed at 20% of excitation power and 40% of MST power. MST traces were analyzed at 1.5 seconds.

For performing experiments with NT-647 labeled Etanercept (Amine Reactive), the concentration of labeled Etanercept was constant (2 nM) while the concentration of rhTNFα was varied between 20 µM –19.1 pM. The assay was performed in PBS containing 0.05% Tween-20. After a short incubation, the samples were loaded into premium glass capillaries. The experiment was performed at 10% of excitation power and 40% of MST power. MST traces were analyzed at 2.5 seconds.

The results were analyzed in the software M.O. Affinity Analysis v.2.3 (NanoTemper Technologies, Germany).

### Size exclusion chromatography-High performance liquid chromatography (SEC-HPLC)

Samples of Adalimumab, Etanercept, rhTNFα and mixtures of them were analyzed using a Jasco system (Japan) with a TSK Gel G4000SW_XL_ column (7.8 mm I.D. x 30 cm) (Tosoh Bioscience, Japan). The chromatographic analyses were performed with a flow rate of 0.5 ml/min and using a 50 mM phosphate-300 mM NaCl buffer (pH 7.5) at room temperature. The detection was performed on a Jasco 2075 Plus UV detector (Jasco, Japan) at 280 nm. The Gel Filtration Calibration Kits (LMW, HMW) (GE Healthcare Life science, Germany) were utilized.

## ACKNOWLEDGMENTS

The authors are thankful to Mr. Manuel J. Iturra and Claudio A. Bastías for their technical support in the preparation and characterization of materials.

## FUNDING AND ADDITIONAL INFORMATION

This work was supported by the Center for Biotechnology and Biomedicine Spa (CBB), and by the ANID (ex-CONICYT) PFCHA/Doctorado Nacional/2014-21141016.

